# 4931414P19Rik: A Chemoattractant Secreted by Neural Progenitors Modulates Microglia Activation and Neuronal Migration During Mammalian Brain Development

**DOI:** 10.1101/2022.12.22.521648

**Authors:** Ivan Mestres, Federico Calegari

## Abstract

Communication between the nervous and immune system is critical for development, homeostasis and response to injury. Prior to the onset of neurogenesis, microglia populate the central nervous system serving as resident immune cells over the course of life. Here, we describe new roles of an uncharacterized transcript upregulated by neurogenic progenitors during mouse corticogenesis: 4931414P19Rik (hereafter named P19). Overexpression of P19 cell-extrinsically inhibited neuronal migration and acted as chemoattractant of microglial cells. Interestingly, effects on neuronal migration were found to result as a direct consequence of P19 secretion by neural progenitors triggering microglia activation and their accumulation within the P19 targeted area. Our findings highlight the critical role of microglia activation during brain development and identify P19 as a novel player in the neuro-immune crosstalk.

## INTRODUCTION

During mammalian brain development, different types of stem and progenitor cells populate the germinal layers including apical and basal progenitors. Apical progenitors within the ventricular zone (VZ) expand through proliferative division, and can also give rise to basal progenitors forming the sub-ventricular zone (SVZ). In turn, basal progenitors generate neurons that migrate through the intermediate zone (IZ) to reach their final destination within the cortical plate (CP) (Lui et al., 2011; Taverna et al., 2014). As a result, the balance between proliferative and neurogenic divisions, and the proper migration of newborn neurons, are fundamental during development to establish the cytoarchitecture and size of the adult brain.

In an attempt to reveal novel mechanisms involved in mammalian corticogenesis, our group previously identified a subset of genes that are transitorily up- or down-regulated specifically by neurogenic progenitors relative to both proliferative progenitors and newborn neurons (Aprea et al., 2013). The functional implication of some of these transcripts identifying the signature of neurogenic commitment, and referred to as up- or down-switch genes, was revealed in subsequent studies focusing on various classes of non-coding RNAs, pioneer transcription factors and epigenetic mechanisms (Aprea et al., 2015; Artegiani et al., 2015; Dori et al., 2019; Dori et al., 2020; Noack et al., 2019). Also among up-switch genes, we found an uncharacterized transcript until now just identified with an annotation number: 4931414P19Rik (human homologue C14orf93; henceforth referred to as P19). To date, we could only find two reports on P19 that associated mutations in its locus with thyroid function and cancer (Liu et al., 2017; Yu et al., 2017). However, the biological implication of this association with thyroid cancer, or underlying molecular mechanism, was not explored. In addition, a role of P19 in tissues other than the thyroid, and particularly the developing brain, was not investigated. This despite the fact that genome-wide association studies linked P19 mutations to schizophrenia (Lam et al., 2019). This prompted us to characterize the function of P19 during mammalian corticogenesis finding its unexpected role as modulator of the crosstalk between neural progenitors and microglia.

While links between the nervous and the immune system are often neglected in the study of neural stem cell fate and neuronal migration, it is intriguing to note that the onset of neurogenesis coincides with microglia colonization of the brain at mouse embryonic day (E) 9-10 (Ginhoux et al., 2010). Specifically, microglia populate the VZ/SVZ and IZ while initially avoiding the CP (Hattori and Miyata, 2018; Hattori et al., 2020), which is colonized only later after neurogenesis is completed (∼E18) (Cunningham et al., 2013). Derived from a precursor in common with microglia, non-parenchymal macrophages also reside within specific niches of the brain under homeostatic conditions including the choroid plexus, meninges and perivascular space (Utz et al., 2020). Finally, upon injury or disease, monocyte-derived macrophages can infiltrate the brain from the blood in a process referred to as neuroinflammation (Han et al., 2021). Importantly, activation of the immune system during gestation is causally linked to several neurodevelopmental disorders (Han et al., 2021). This raises several fundamental questions pertaining to which signals attract brain microglia and/or macrophages to populate specific layers of the cortex and whether their activation plays any role in neural cell fate specification and/or migration of newborn neurons. While some attempts were made to address the role of microglia in corticogenesis (Arnò et al., 2014; Cunningham et al., 2013; Hattori and Miyata, 2018; Hattori et al., 2020), the answer to these questions remains elusive.

Here, we identify P19 as a novel cell-extrinsic chemoattractant of microglia promoting their activation and accumulation within the germinal zones. In turn, we show that P19 secretion by neural progenitors is critical in mediating the neuro-immune crosstalk and regulate neuronal migration during brain development.

## RESULTS

### P19 cell-extrinsically controls neuronal migration

Our group previously identified P19 as an up-switch gene, i.e.: a transcript identifying the signature of neurogenic commitment whose expression during corticogenesis is the highest in progenitors undergoing neurogenic division relative to both proliferative progenitors and newborn neurons (Aprea et al., 2013) (Supp. Fig. 1A and B). Specifically, P19 was not only among the top 50% most expressed transcripts in the embryonic brain but also, when analysing proliferative versus neurogenic progenitors and neurons of the E14 mouse cortical wall, it was upregulated ca. 2-fold in neurogenic progenitors relative to the other two cell types (normalized reads: 241 ± 15, 392 ± 17, and 242 ± 7, respectively; p < 0.05). To corroborate our previous findings of the E14 mouse cortex, we took advantage of single-cell transcriptome analyses (Cao et al., 2019; Telley et al., 2019) confirming a ca. 1.5-fold higher expression of P19 in neurogenic progenitors compared to radial glial cells and postmitotic neurons when evaluated between E9 and E13. Moreover, by combining fluorescent immunohistochemistry with in situ hybridization in E14 brain sections, we additionally validated that Tbr2+ neurogenic progenitors within the SVZ displayed nearly 3-fold higher levels of P19 expression relative to both proliferating progenitors and neurons (Supp. Fig. 1C and D).

To investigate a putative function of P19, we next performed in utero electroporation of the cortical wall at E13 using a plasmid encoding a red fluorescent protein (RFP) alone as control, or together with P19 under an independent constitutive promoter. Two days later, brains were collected to evaluate effects on corticogenesis by immunohistochemistry. At E15, we observed that the largest fraction of RFP+ cells electroporated with either control or P19 plasmids were retained within the proliferative areas (VZ/SVZ) or migrated into the IZ (Fig. 1A). While no major difference between control and P19 electroporated brains was observed in the proportion of RFP+ cells within the VZ/SVZ or IZ, P19 electroporated brains displayed an almost complete lack of RFP+ cells in the CP relative to controls (CP RFP = 17.3 ± 2.2% vs. P19 = 2.5 ± 0.5%; p < 0.05) (Fig. 1A and B). This effect was unlikely to be due to apoptosis since immunolabeling with caspase 3 showed no difference between brains electroporated with either construct (Supp. Fig. 1E and F).

**Figure 1.**
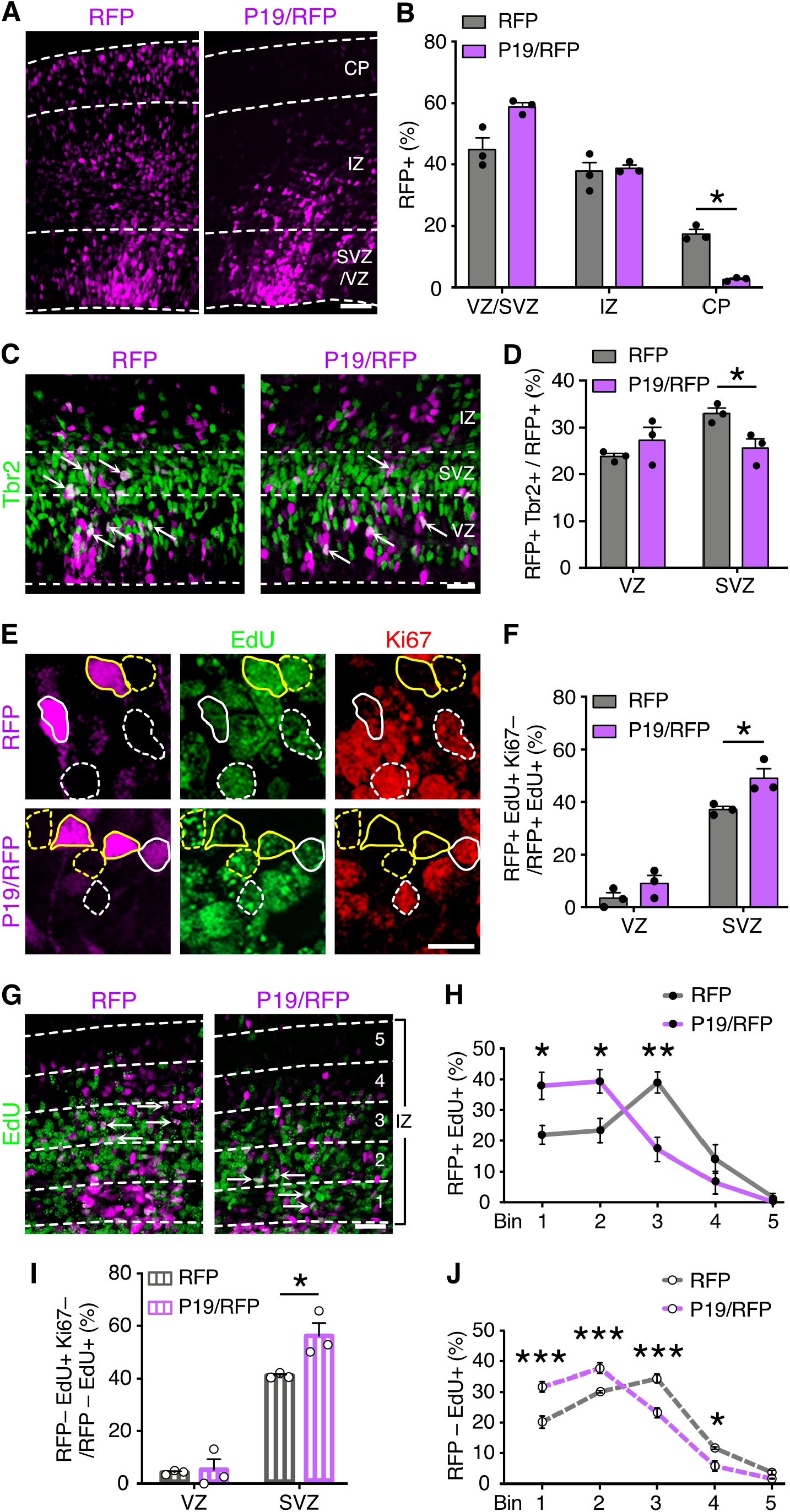
P19 is an up-switch gene involved in corticogenesis. (A, C, E, G) Fluorescence images of coronal sections of E15 brains two days after electroporation with control RFP or P19/RFP plasmids (pseudocolored in magenta) and immunolabeled with markers of basal progenitors, proliferation and S-phase (Tbr2, Ki67 and EdU, respectively, as indicated). Dashed lines delimit the ventricular/subventricular zones (VZ/SVZ), intermediate zone (IZ) and cortical plate (CP) (A and C), or bins within the IZ (G). Contour lines in (E) indicate examples of cells positive (continuous) or negative (dashed) for RFP; and Ki67+ or Ki67– cells (white and yellow lines, respectively). Lower magnification panels of (E) are shown in (Supp. Fig. 1K). (B, D, F, H-J) indicate percentages of cells quantified from the respective panels (A, C, E and G) and considering RFP+ (B, D, F and H) or adjacent RFP– (I and J) cells. Quantifications are depicted as bar graphs with individual values (B, D, F, I) or stacked lines (H, J) ± SEM. Arrows in C and G point to double positive cells. Two-way ANOVA and Bonferroni’s post-hoc test were used to assess significance (* p < 0.05, ** p < 0.01, *** p < 0.001). Scale bars = 50 µm (A, G), 25 µm (C), 10 µm (E).

We next sought to examine whether P19 modulated different aspects of neural progenitor cell fate. We started by evaluating the mitotic index within the VZ or SVZ by quantifying the proportion of RFP+ transfected cells counterstained with the mitotic marker phospho-histone 3 (PH3), which revealed neither major nor significant differences between control or P19 vectors (Supp. Fig. 1G and H). In contrast, we observed a minor, although significant, decrease in the proportion of electroporated cells positive for the basal progenitors marker Tbr2 within the SVZ, but not the VZ, of P19 electroporated brains relative to control (SVZ RFP = 33.0 ± 0.9% vs. P19 = 25.6 ± 1.6%; p = 0.04) (Fig. 1C and D). Additionally, we performed electroporations in the *Btg2*::GFP reporter mouse line as a mean to directly identify progenitors (either apical or basal) committed to neurogenic divisions (Haubensak et al., 2004). This experiment showed that compared to brains transfected with control plasmids, P19 again triggered a decrease in the proportion of *Btg2*::GFP+ progenitors specifically within the SVZ, but not the VZ, whose magnitude was similar to the one assessed by Tbr2 (SVZ RFP = 29.0 ± 1.5% vs. = 21.6 ± 1.4%; p = 0.04) (Supp. Fig. 1I and J).

The almost complete absence of neurons in the CP upon P19 overexpression was hard to solely explain by a decrease in neurogenic divisions given that the observed reduction in Tbr2+ (basal) and Btg2+ (neurogenic) progenitors within the SVZ was minor and barely significant. Alternatively, we argued that a more likely explanation for the lack of RFP+ cells in the CP was that P19 impaired the migration of newborn neurons, which would also result in an increase in the proportion of Tbr2– and Btg2– postmitotic neurons retained within the SVZ. We reasoned that this was even more likely considering that while the proportion of RFP+ cells in the IZ of P19 electroporated brains was similar to control, their distribution was clearly biased apically towards the SVZ (Fig. 1A).

To validate a possible increase in postmitotic neurons within the SVZ upon P19 overexpression, we assessed cell cycle exit by means of a single injection of EdU 24 h after electroporation and collecting the brains 24 h thereafter. Immunolabeling with the proliferation marker Ki67 was performed to assess cells that exited the cell cycle within this developmental time but that were still retained within the SVZ. Interestingly, we found not only that an increased proportion of RFP+ cells exited the cell cycle (EdU+ Ki67–/EdU+) in the SVZ of P19 electroporated brains but also that this increase matched in magnitude (SVZ RFP = 37.0 ± 0.9% vs. P19 = 49.0 ± 2.9%; p = 0.04) (Fig. 1E and F) the decrease in Tbr2+ or Btg2+ cells described above.

Next, we sought to directly confirm the effects of P19 overexpression on neuronal migration. To this aim, experiments in which mice were administered EdU 24 h before sacrifice (see above) were used as a birth-dating strategy to assess the migration of RFP+ EdU+ cells by evaluating their distribution across 5 equally-sized bins within the IZ. This analysis revealed that upon electroporation with control plasmids twice as many RFP+ neurons localized halfway through the IZ (bin 3) relative to any other bin. In contrast, the majority of P19-transfected cells accumulated nearer the SVZ and were almost exclusively limited to bins 1 and 2 (Fig. 1G and H).

Together, our analyses revealed that the physiological expression of the up-switch gene P19, as previously identified by our group (Aprea et al., 2013), is important for the proper migration of newborn neurons with little, if any, effect in regulating the balance between proliferative and neurogenic divisions.

While extending previous functional characterizations of up- and down-switch genes by our group (Aprea et al., 2015; Artegiani et al., 2015; Dori et al., 2019; Dori et al., 2020; Noack et al., 2019), in the current analysis of P19 we observed a completely unexpected phenomenon. To our surprise, when performing the analyses described above not only among RFP+ targeted cells but also among their neighbouring RFP–, untransfected cells, the same effects were found. This included both an increased cell cycle exit within the SVZ (Fig. 1I) and decreased neuronal migration (Fig. 1J). Even more, we noted that the magnitude of these effects was virtually identical when assessed among RFP+ and RFP– cells of the same brains (Fig. 1F vs. I and H vs. J). In essence, these results indicated that the effects of P19 were cell-extrinsic.

### P19 is a secreted chemoattractant of microglia

Given the surprising cell-extrinsic effects of P19, we next sought to inspect features within its sequence that would provide indications about its function and secretion.

However, neither an analysis of the primary amino acid sequence by InterPro (Blum et al., 2021) nor secondary structure by HHPred (Zimmermann et al., 2018) revealed any conserved catalytic domain that could be used to deduce a molecular function. Nevertheless, two important aspects of P19 were highlighted. First, predictions by NLS Mapper (Kosugi et al., 2009) and NLStradamus (Nguyen Ba et al., 2009) identified two bipartite nuclear localization signals starting at residues 291 and 369 (Fig. 2A; Supp. Table 1). Second, other four bioinformatic prediction software (TOPCONS, Phobius, PrediSi, and SignalP) (Hiller et al., 2004; Käll et al., 2004; Petersen et al., 2011; Tsirigos et al., 2015) predicted a signal peptide within the first 17 amino acids from the N-terminal and a cleavage site between amino acids 17 and 18; both of which are hallmarks of secreted proteins (Fig. 2A; Supp. Table 1). The rest of P19 sequence was predicted to just be “non-cytoplasmic”. We reasoned that while the significance of the predicted nuclear localization signals remained unclear, the presence of an N-terminal signal peptide, cleavage site, lack of predicted transmembrane domains, and “non-cytoplasmic” sequence were all consistent with P19 being a soluble protein secreted via vesicular exocytosis.

**Figure 2.**
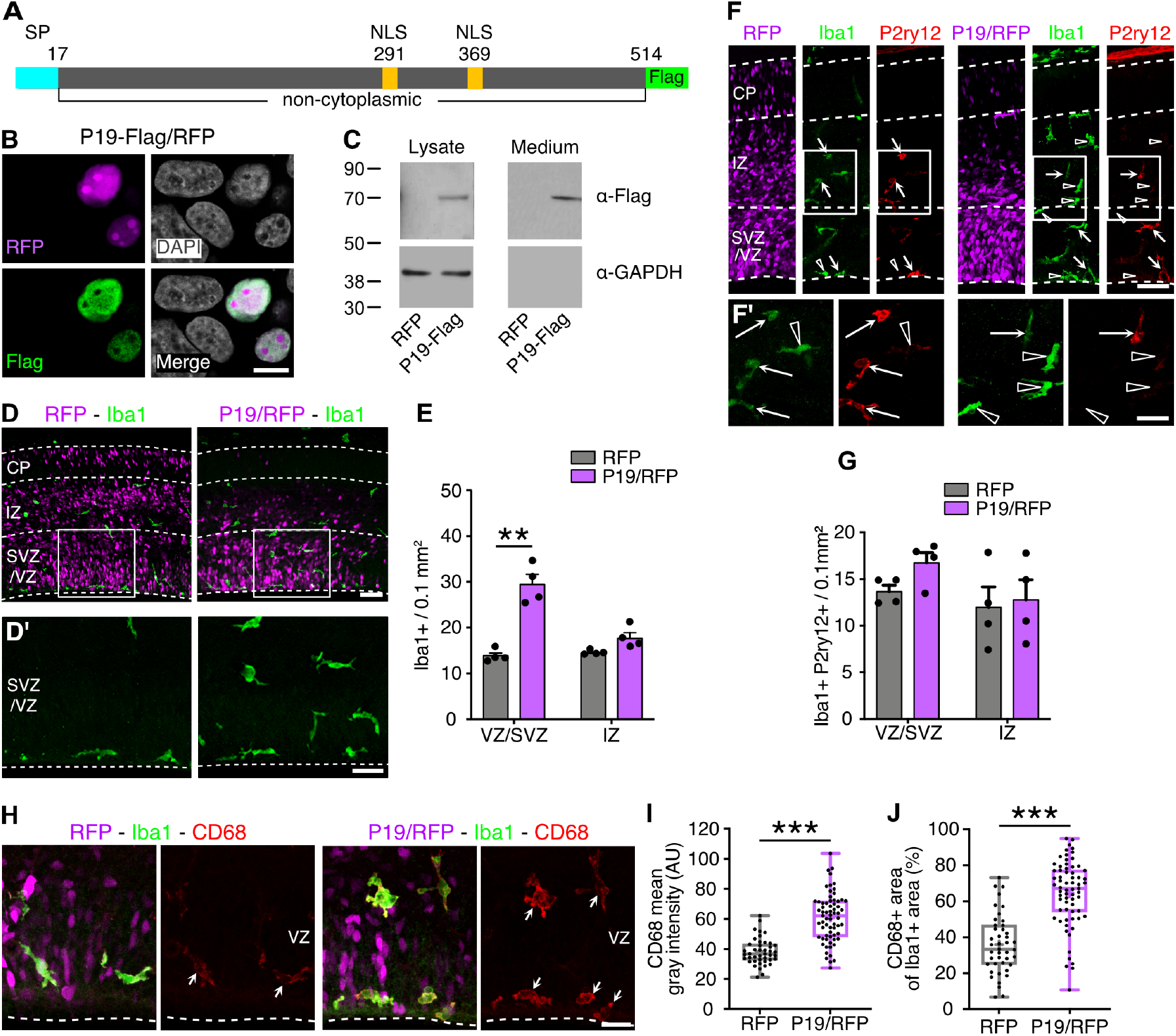
P19 is a secreted chemoattractant of microglia. (A) Scheme of P19 primary amino acid sequence, including the predicted signal peptide (SP, light blue) and nuclear localizing signals (NLS, yellow). (B) Fluorescence picture of HEK293cells transfected with P19-Flag and RFP and counterstained as indicated. (C) Western blots of cell lysates or culture medium of HEK293cells transfected with RFP or P19-Flag upon anti-Flag or GAPDH detection. (D, F and H) Fluorescence pictures of coronal brain sections 48 h after electroporation counterstained with Iba1 (green) and the resident microglia marker P2ry12 (red, F) or CD68 (red, H). Insets in (D and F) are magnified in (D’ and F’). Arrows point to Iba1+ and P2ry12+ or CD68+ cells, while arrowheads show cells positive only for Iba1. (E, G) Quantification of cell density for the respective markers. (I, J) Area (I) or mean gray intensity levels (J) of CD68 within Iba1+ cells. Quantifications are depicted as bar graphs ± SEM (E, G) or box-plots (I, J). Two-way ANOVA and Bonferroni’s post-hoc test (E, G) or two-tailed unpaired t test (I, J) were used to assess significance (** p < 0.01, *** p < 0.001). Scale bars = 10 µm (B), 50 µm (D, F), 25 µm (D’), 20 µm (H).

Next, to confirm the nuclear localization and secretion of P19, we used HEK293 cells transfected with a dual promoter construct and independently co-overexpressing i) P19 fused to a Flag tag at the C-terminus and ii) a nuclear localized RFP (P19-Flag/RFP). Subsequent Flag immunolabeling showed that P19 localized to the nucleus (with the exception of nucleoli) (Fig. 2B). To validate whether P19 was also secreted, a similar experiment was performed but this time obtaining cell lysates and culture media to assess P19-Flag by western blot analyses. This revealed a single band at 70 kDa in both the cellular and medium fractions (Fig. 2C). The increase between the predicted (58 kDa) and the observed size can be explained by possible post-translational modifications, such as glycosylation, which is typical of secreted proteins. Consistent with bioinformatic predictions, these experiments supported both P19 localization to the nucleus and its secretion.

Finding that P19 is a secreted protein opened up the possibility that its cell-extrinsic effects upon in utero electroporation were not limited to neural progenitors and newborn neurons but could also apply to any other cell type of the developing cortex. While astrocytes and oligodendrocytes first appear at later stages of development (Rowitch and Kriegstein, 2010), at the time of P19 overexpression (E13-15), and specifically within the cortical wall, there are essentially only three cell types in addition to neural progenitors and projection neurons, namely interneurons, endothelial cells, and macrophages, with the latter including microglia proper, perivascular macrophages or, potentially, infiltrating monocytes. Hence, to test the effects of P19 overexpression on these cell types, we subjected brain slices electroporated as described above to immunolabeling with antibodies against calbindin (interneurons), CD31 (endothelial cells) and Iba1 (microglia/macrophages).

When analysing the migratory stream of interneurons that was the closest to the SVZ (Marín et al., 2010), where most electroporated cells localize, we could not find any obvious change in the total number of calbindin+ cells per area, nor in their distribution across 3 equally-sized bins perpendicular to the ventricular surface (to account for tangential, rather than radial, migration of this neuronal population) (Supp. Fig. 2A-C). Similarly, when assessing the blood vessels by means of CD31 labelling none of the parameters considered, including their density, length, diameter and branching, were changed upon P19 overexpression (Supp. Fig. 2D-I). Finally, we assessed brain macrophages by immunolabeling with the marker Iba1. This showed, in agreement with previous reports (Cunningham et al., 2013), that Iba1+ cells almost exclusively localized within the VZ/SVZ and IZ (Fig. 2D) and that their distribution was homogenous within and outside the RFP+ electroporated area of control brains. In contrast, remarkably, microglia displayed a greater density within the electroporated area of P19 targeted brains (Fig. 2D and E).

Specifically, in P19 electroporated brains, and particularly within the VZ/SVZ, the number of Iba1+ cells more than doubled when compared to brains electroporated with control plasmids (VZ/SVZ RFP = 13.9 ± 1.2 vs. P19 = 29.4 ± 4.0 Iba1+ cells per 0.1 mm^2^; p < 0.01). Clearly, macrophages themselves were not targeted by electroporation given that these cells do not contact the ventricular surface where plasmids were injected and, consistently, no colocalization of RFP with Iba1 was found (e.g. Fig. 2D). In sum, this implied that macrophages accumulation within the targeted area resulted from P19 secretion from neural progenitors.

Given that Iba1 does not discriminate between different classes of macrophages, we next sought to identify distinct cell sub-populations by labelling with a combination of molecular markers, namely: Ccr2, CD206 and Lyve1 for non-parenchymal or infiltrating macrophages and P2ry12 for microglia. We noticed both in control and P19 electroporated brains that Ccr2+ macrophages were negligible within the brain areas analysed and that the few detected were negative for Iba1 (Supp. Fig. 2J). In addition, the use of CD206 and Lyve1 revealed non-parenchymal macrophages within the choroid plexus and meninges but not at the level of the perivascular space (data not shown), which is consistent with reports showing the negligible contribution by this population at the stages of development analysed (Utz et al., 2020). Altogether these results excluded the possibility that accumulation of Iba1+ cells upon P19 overexpression resulted from infiltrating monocytes or perivascular macrophages colonizing the brain parenchyma.

Intriguingly, while in control RFP electroporated brains most Iba1+ cells were also immunoreactive for the canonical microglial protein P2ry12, only a fraction of Iba1+ cells were labelled with P2ry12 upon P19 overexpression (Fig. 2F and G). The downregulation of P2ry12 suggested the possibility that P19 triggered microglia activation (Haynes et al., 2006). To validate this, we used the lysosomal marker CD68, which is upregulated in reactive microglia (Papageorgiou et al., 2016). Quantification of CD68 signal intensity and area occupied within Iba1+ cells was almost doubled after P19 overexpression compared to control RFP electroporations (CD68 mean gray intensity RFP = 38.2 ± 1.3 vs P19/RFP = 60.8 ± 1.9 AU, p < 0.0001; CD68+ area RFP = 35.9 ± 2.4 vs P19/RFP = 64.5 + 2.2%, p < 0.0001) (Fig. 2H-J). These observations further support the notion that P19 secretion from neuronal progenitors triggers microglia activation and their accumulation within the targeted area.

So far, our observations were based on overexpression data. Thus, we next attempted the converse manipulation of P19 knockdown. However, in utero electroporation of a shRNA plasmid, validated in HEK293 cells to silence P19 by ca 70%, did not result in any change in the parameters described above (data not shown). These negative results are not surprising given that electroporation targets only 25-30% within a background of unmanipulated cells that still express and secrete P19. In a second attempt, we electroporated endoribonuclease-digested small interfering (esi) RNAs triggering RNAi homogeneously within the tissue (Calegari et al., 2002). But in this case an increase in Iba+ cells was observed not only when targeting P19 but also an unspecific sequence (Luciferase; data not shown). This result suggests that caution is needed when interpreting data derived from esiRNAs gene silencing in the developing brain due to a potential unspecific activation of immune response. While neither of the two RNAi methods provided conclusive results, we continued our study focusing on overexpression deferring to future studies the generation of P19 knock-out mice.

### P19 increases microglia speed and migration from neighbouring areas

Intrigued by our findings, and knowing that embryonic microglia react quickly upon local environmental changes (Prinz et al., 2019), we investigated whether microglia accumulation upon P19 overexpression was also detected 24 h, rather than 48 h, after electroporation, which is the minimum time necessary to reliably identify the RFP+ targeted area. In fact, a greater density of microglia was observed also at this shorter timepoint, with twice as many Iba1+ cells in the RFP+, P19 overexpressing area relative to the adjacent RFP– area of the same brains (Fig. 3A and B). We found this increase in microglia accumulation already 24 h after electroporation remarkable considering that several hours are necessary for P19 to be expressed, translated, secreted and accumulate in the extracellular space to the levels needed to trigger a microglia response.

**Figure 3.**
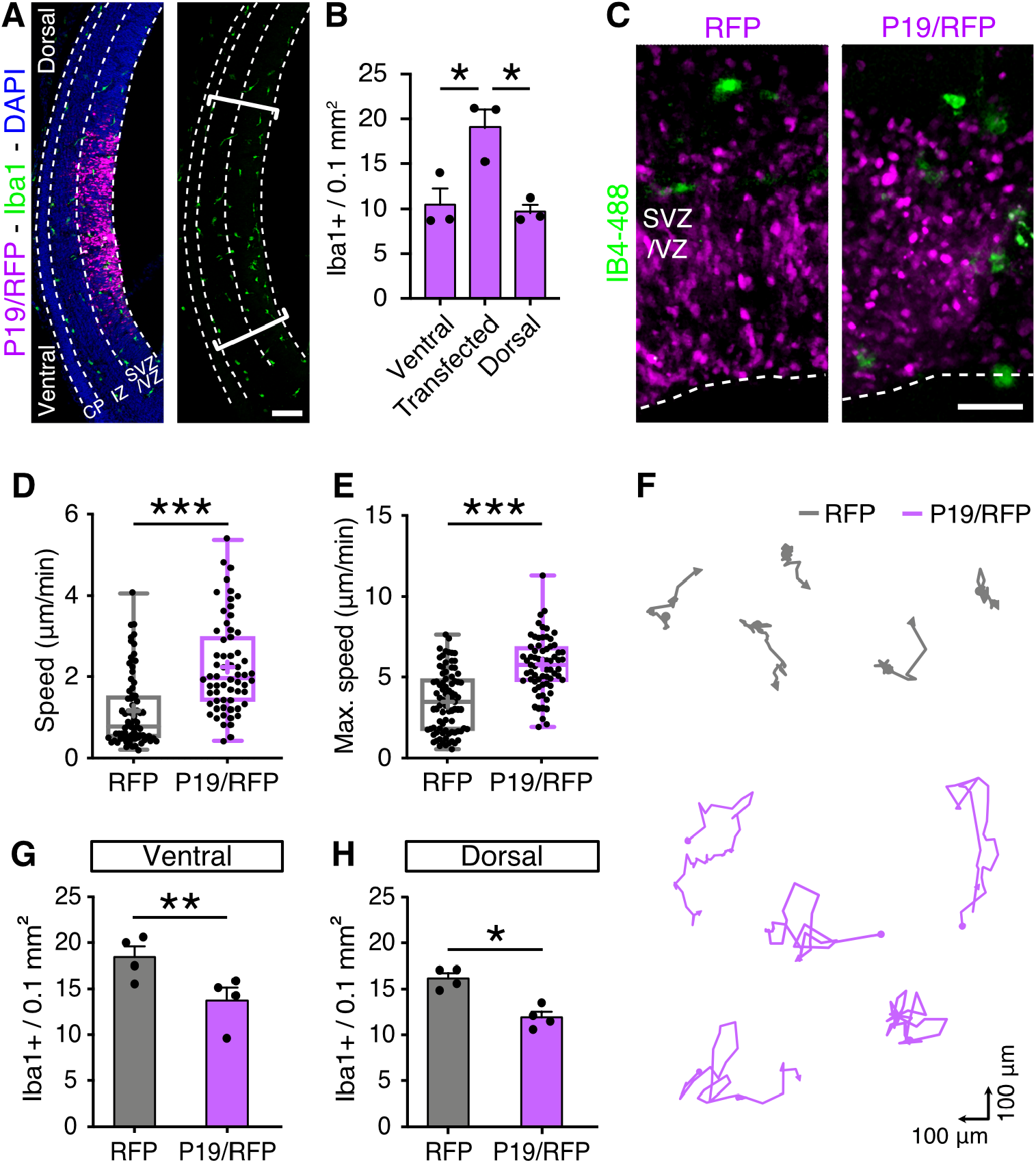
P19 increases microglia migration. (A) Coronal brain section 24 h after electroporation counterstained with Iba1; brackets delimit the electroporated area. (B) Quantification of Iba1+ cell density. (C-F) Fluorescence pictures (C), quantifications (D and E) and representative trajectories (F) of IB4-488 labelled microglia (green) obtained by time-lapse microscopy to calculate average and maximum speed (as indicated). (G, H) Iba+ cell density at adjacent RFP– areas ventrally (G); or dorsally (H) to the electroporated areas. Quantifications are depicted as box plots (D, E) or bar graphs ± SEM (B, G, H). One-way ANOVA and Tukey’s post-hoc test (B), or two-tailed unpaired t test (D, E, G, H) were used to assess significance (* p < 0.05, ** p < 0.01, *** p < 0.001). Scale bars = 100 µm (A), 50 µm (C).

Next, we investigated whether P19 overexpression in neural progenitors triggered changes in other key features characterizing microglia by, specifically, assessing their i) morphology, ii) proliferation as well as iii) phagocytosis and iv) migratory behaviour.

First, throughout prenatal development microglia transition from an amoeboid to a branched morphology. Amoeboid microglia present fewer processes and cellular extensions that are shorter than their soma while, conversely, branched microglia exhibit longer cellular processes and often ramified (Swinnen et al., 2013). Such coexistence of amoeboid and branched microglia was reflected in brain sections upon electroporation but with no change in their proportions between control or P19 targeted brains (Supp. Fig. 3A and B).

Second to investigate if accumulation of microglia upon P19 overexpression was due to their increased proliferation, we treated mice with a single injection of EdU one day after electroporation at E13 and harvested their brains 24 h thereafter at E15. Similar to previous reports (Arnò et al., 2014; Swinnen et al., 2013), about 25% of Iba1+ microglia were found proliferating (EdU+) but, again, no difference was found between control and P19 targeted brains (Supp. Fig. 3C and D).

Third, microglia can phagocytose neural progenitor cells during development (Cunningham et al., 2013) and display a highly dynamic migratory behaviour (Hattori and Miyata, 2018; Swinnen et al., 2013). To assess both parameters, organotypic slice cultures were prepared 24 h after in utero electroporation at E13 and treated with isolectin-B4 coupled to Alexa flour 488 allowing direct visualization of microglia (Dailey et al., 2013) (Fig. 3C). Next, dual colour time-lapse microscopy was performed over the course of additional 12 h to evaluate the interaction between transfected progenitors (RFP+) and microglia (Alexa-488+). Similarly to previous studies (Cunningham et al., 2013), phagocytosis events were scored when red and green signals colocalized for more than 45 min and followed by fragmentation of the RFP signal. By this approach, no difference was observed in phagocytosis by microglia between brains electroporated with control or P19 vectors (RFP = 6.75 ± 1.4 vs. P19 = 8.37 ± 1.7 phagocytosis events in 12 h; p = 0.48, two-tailed unpaired t test).

Finally, fourth, having excluded effects of P19 overexpression on microglia morphology, proliferation and phagocytosis, the same time-lapse imaging was used to assess their migration. While in control targeted slices, comparable to previous reports (Swinnen et al., 2013), microglia migrated at a speed of 1.16 ± 0.1 µm/min, remarkably, microglia were twice as fast, 2.23 ± 0.1 µm/min (p < 0.001), in the targeted area of P19 electroporated brains (Fig. 3D). This faster migration of microglia did not seem to result from shorter pause intervals between displacements (defined as the time spent at a speed lower than 10% of the mean of control) (pause intervals RFP = 25.6 ± 3.8 min vs. P19 = 17.6 ± 4.9 min; p = 0.2, two-tailed unpaired t test). Rather, microglia increased average speed upon P19 overexpression resulted from a higher maximal speed that almost doubled from 3.50 ± 0.2 to 5.73 ± 0.2 µm/min (p < 0.001) (Fig. 3E). Consistent with a higher migration speed, plotting individual cell trajectories revealed that microglia covered a larger surveillance area in P19 transfected brains compared to control (Fig. 3F). Our time-lapse imaging together with our previous quantification of EdU incorporation suggested that active microglia accumulated into the transfected area by migration from neighbouring regions rather than by local proliferation. To verify this, we evaluated adjacent RFP– areas, both ventrally and dorsally to the electroporated site and indeed found a reduced number of Iba1+ cells compared to similar areas of control RFP targeted brains (Fig. 3G and H).

Altogether, these results showed that P19 functions as an activator and chemoattractant of microglia and that its expression and secretion by neural progenitors results in their accumulation by means of an increased migration.

### Microglia accumulation and impaired neuronal migration are causally linked

So far, our analyses showed that P19 i) is primarily expressed by basal neurogenic progenitors of the VZ/SVZ, ii) is a secreted molecule whose overexpression cell-extrinsically impairs neuronal migration, and iii) act as a chemoattractant of microglia promoting their activation and migratory behaviour. Together, these observations raise several important questions: first, whether P19 secretion per se is a direct cause of the phenotypes observed or, alternatively, these phenotypes arise from secondary effects that electroporated progenitors trigger in response to their P19 overexpression. Second, since the CP is devoid of microglia at the stages of development being considered, whether P19 could anticipate colonization of this layer when overexpression was performed at the level of the pial, rather than apical, boundary of the cortical wall. And third, whether the observed impairments in neuronal migration and accumulation of microglia are causally linked rather than being two independent effects.

To address the first two questions, we sought to provide the developing cortex with an ectopic source of P19 but this time using a heterologous cell type instead of endogenous neural progenitors. To this aim, HEK293 cells were transfected with a control or P19 overexpressing plasmid and embedded into a collagen drop. Adapting our use of ex vivo slice cultures, the HEK293-containing collagen drop was then placed adjacent to the pial surface of E14 brain slices in a similar medio-lateral location as the one usually targeted by electroporation (Fig. 4A). After 24 h of culture, we evaluated the number of microglia by immunolabeling for Iba1 and found that, similarly to experiments in vivo (Fig. 2D), the proximity to P19-transfected cells triggered an increased density of microglia in the IZ relative to that observed when using HEK293 cells transfected with control vectors (Fig. 4B and C) (IZ RFP = 5.3 ± 0.2 vs. P19 = 16.3 ± 2.9 cells per 0.1 mm^2^; p < 0.01). Notably, however, since this time the source of P19 was from the pial surface, accumulation of microglia was observed at the basal boundary of the IZ instead of its apical boundary with the SVZ (Fig. 4B). This implies not only that microglia accumulation is independent from the cell type releasing P19 but also from the topological source of its origin. Despite of this, we observed that the higher density in microglia within the IZ did not result in their increased invasion of the CP (Fig. 4C) suggesting that P19 alone is not potent enough to override the mechanisms preventing microglia colonization of this cortical layer at this developmental stage. Additionally, while some microglia are known to arise in the CP of cultured ex vivo brain slices (Hattori et al., 2020; Swinnen et al., 2013), that are never observed in physiological conditions, the abundance of these microglia was not influenced by P19.

**Figure 4.**
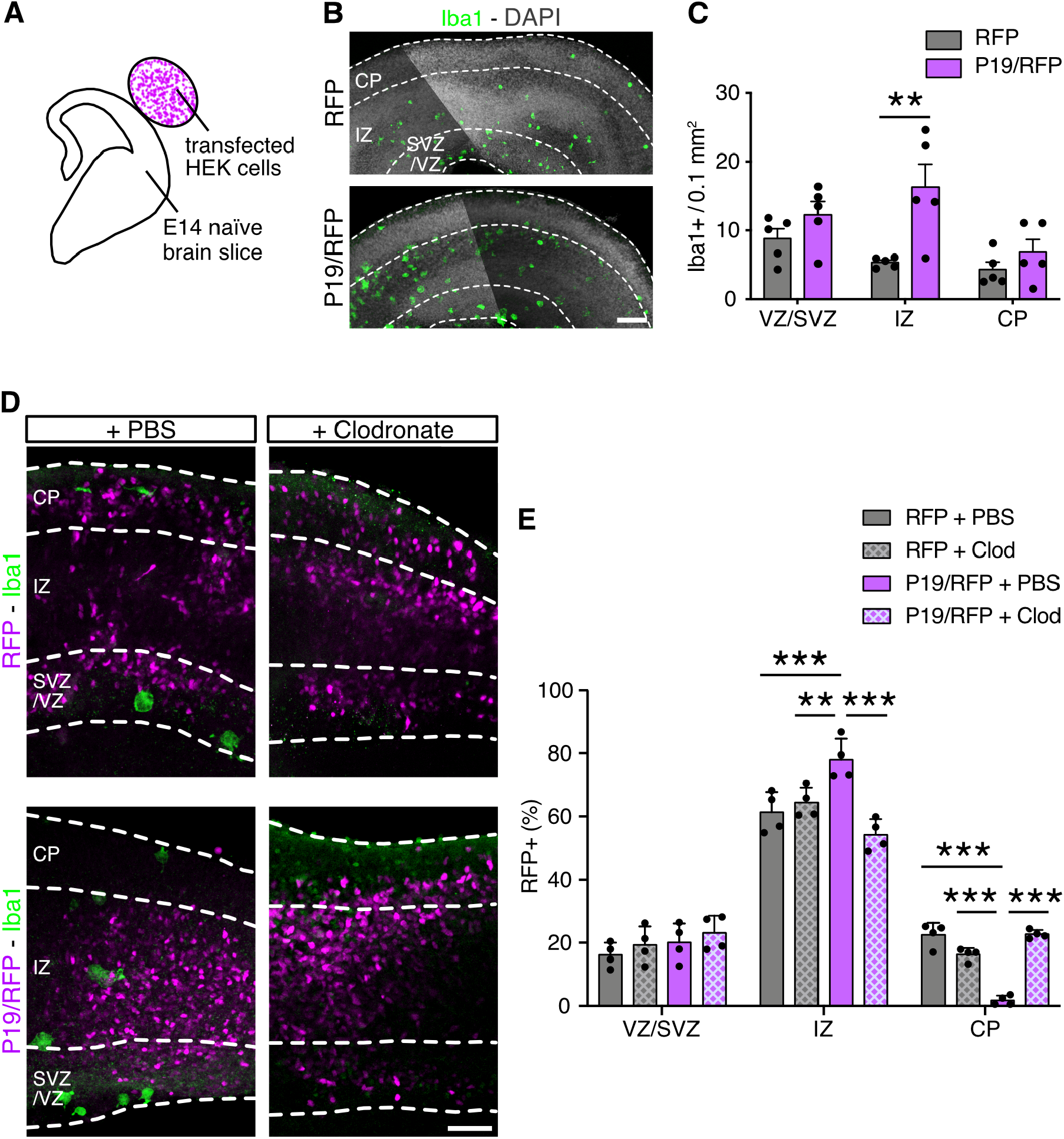
Depletion of microglia rescues P19-triggered impairment in neuronal migration. (A) Drawing of brain slice co-culture with transfected HEK293cells embedded in a collagen drop. (B) Fluorescence images of brain slices exposed to HEK293cells transfected either with RFP or P19 as depicted in (A) followed by immunolabeling and quantification (C) of Iba1+ microglia. (D and E) Fluorescent images and quantifications of electroporated brain slices incubated with liposomes containing either PBS or Clodronate. Quantifications are depicted as bar graphs with individual values ± SEM. Two-way ANOVA and Bonferroni’s post-hoc test were used to assess significance (** p < 0.01, *** p < 0.001). Scale bars = 100 µm (B), 50 µm (D).

Finally, third, we investigated whether P19-dependent accumulation of microglia was necessary to inhibit neuronal migration. To this aim, P19 was overexpressed as described above, but this time microglia accumulation was prevented by exposing brain slices to liposomes loaded with clodronate, which are selectively engulfed by microglia, inducing their cell-specific death (Kumamaru et al., 2012). When assessing slices exposed to PBS-loaded liposomes, and recapitulating previous results in vivo (Fig. 1A), we noticed that compared to electroporation with control plasmids, P19-targeted slices contained fewer RFP+ cells in the CP (CP RFP + PBS = 22.5 ± 1.6 vs. P19 + PBS = 1.7 ± 0.6%; p < 0.001) that instead accumulated in the IZ (Fig. 4D and E). In contrast, when assessing the distribution of RFP+ cells in slices treated with clodronate-loaded liposomes, the P19-driven impairment in neuronal migration was not observed and the distribution of RFP+ cells was undistinguishable from control electroporations (Fig. 4D and E) (CP RFP + clodronate = 16.3 ± 0.8% vs. P19 + clodronate = 22.8 ± 4.2%; p = 0.38). In essence, ablation of microglia completely rescued the P19-triggered effect on neuronal migration. In turn, this suggested that while ablation of microglia in a control background was not sufficient to hinder neuronal migration, the converse accumulation of microglia upon P19 overexpression was necessary to impair neuronal migration.

### Delayed migration triggered defects in cortical layering

To evaluate if the phenotypes observed upon P19 overexpression persisted at later stages of development we decided to analyse the brains at E18, i.e. five days, instead of two, after electroporation at E13. We observed that, compared to the earlier time-point analysed (E15), P19 overexpression increased the magnitude of cells accumulated within the germinal zones (VZ/SVZ RFP = 5.4 ± 0.5% vs. P19 = 18.7 ± 1.1%; p = 0.0015) at the expense of their migration into the CP (RFP = 79.6 ± 1.9% vs. P19 = 63.5 ± 3.1%; p = 0.0001) (Fig. 5A and B). Similarly, microglia density remained higher in the VZ/SVZ of P19 electroporated brain slices (RFP = 40.2 ± 3.8 vs. P19 = 54.3 ± 1.6 Iba1+ cells per 0.1 mm^2^; p < 0.005) (Fig. 5A and C); although to a lower degree compared to E15 brains, which is consistent with the fact that electroporation is a transient overexpression system. At this time-point (E18), the cortex starts to acquire a layered structure, and neurons define their subtype identity. In agreement with the finding that early-born neuron migration is delayed, fewer P19+ neurons were immunolabeled with the deep-layer marker Ctip2 (Fig. 5D and E). Altogether, these results suggest that while the effects on microglia activation and accumulation was attenuated over time, their influence on neuronal migration, and accordingly cellular identity and cortical layering, was not compensated at later stages of development.

**Figure 5.**
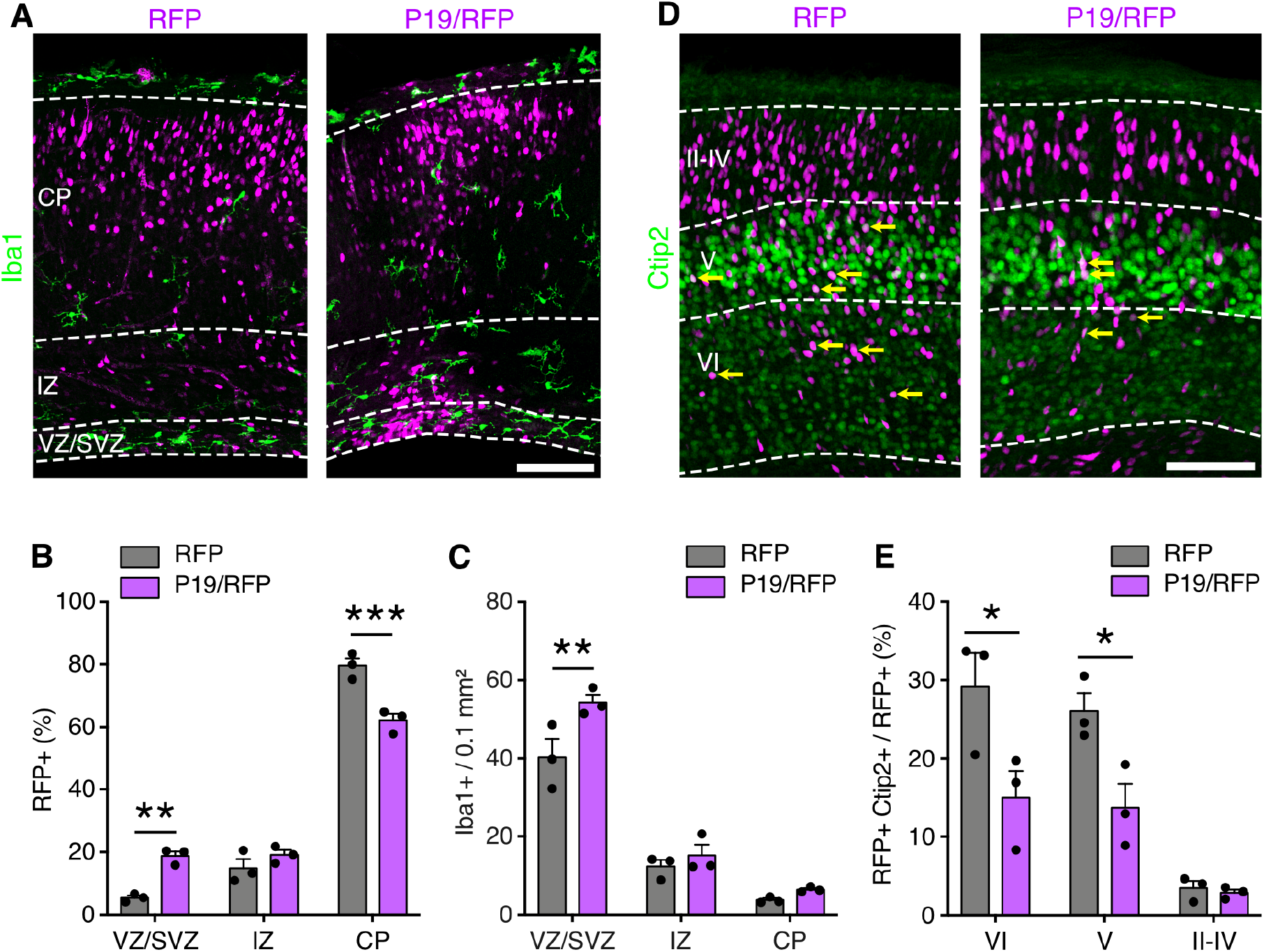
Delayed migration after P19 overexpression impaired cortical layering. (A and D) Fluorescence images of brain slices five days after electroporation at E13 counterstained with Iba1 (A) or Ctip2 (D). Yellow arrows point to RFP+ and Ctip2+ cells. Bar graphs indicate percentage of transfected cells (B), density of Iba1+ cells (C), or percentage of Ctip2+ cells (D). Quantifications are depicted with individual values ± SEM. Two-way ANOVA and Bonferroni’s post-hoc test were used to assess significance (* p< 0.05, ** p < 0.01, *** p < 0.001). Scale bars = 100 µm (A, D).

## DISCUSSION

Microglia, non-parenchymal macrophages and infiltrating monocytes were traditionally studied in the context of the adult nervous system where they engulf apoptotic cells or pathogens and modulate the inflammatory response (Prinz et al., 2019). However, the distinct function of these three macrophage cell types remains elusive (Utz et al., 2020). More recently, tissue resident microglia were also suggested to execute more sophisticated functions including to regulate synapse remodelling and plasticity and, as a result, cognitive performance (Favuzzi et al., 2021; Nguyen et al., 2020; Roy et al., 2022). In contrast, a role of microglia in controlling neural stem cell fate and neuronal migration during brain development has received much less attention. In this work, we provide the first description of the role of P19 in brain development and in doing so reveal novel insights into the neuro-immune crosstalk during corticogenesis. Our results point to P19 as a novel factor secreted by neural progenitors acting as chemoattractant and activator of microglia and ultimately influencing neuronal migration.

Intriguingly, in our study P19 secretion selectively acted at the level of microglia without any evident effect on other types of macrophages including non-parenchymal and infiltrating monocytes. The nature of this selectivity remains unclear. On the one hand, perivascular macrophages within the P19 targeted area of the cortex are few at the stages of development analyzed (Utz et al., 2020) raising the possibility that their accumulation could not be revealed by our study. On the other hand, infiltration of monocytes and their maturation into macrophages is a relative slow process requiring several days (Ajami et al., 2011) and difficult to observe within the few days following electroporation. However, the possibility remains for P19 to be a microglia-specific chemoattractant rising questions about the significance of its up regulation and secretion specifically by neurogenic progenitors within the SVZ.

Few studies have addressed the mechanisms instructing microglia to colonize the central nervous system during development. Intriguingly, microglia colonization and neurogenic commitment closely coincide during corticogenesis (Ginhoux et al., 2010) raising the possibility that factors released by neurogenic progenitors may act as chemoattractants of microglia guiding their migration into the brain parenchyma. Consistent with this hypothesis, at least one factor was previously reported to be secreted by neural progenitors and attract microglia: Cxcl12 (C-X-C motif chemokine ligand 12) (Arnò et al., 2014). Here, we report the second such factor: P19, that selectively activates and promotes the migration of microglia. Notably, while inspecting our previous catalogue of switch genes characterizing neurogenic commitment (Aprea et al., 2013), we found that both Cxcl12 and P19 are on-switch genes. Such common feature is likely not coincidental considering that on-switch genes are an underrepresented class of transcripts consisting of less that 1% of the transcriptome of neurogenic progenitors (Aprea et al., 2013). While not sufficient to override the signals preventing the colonization of the CP, it is tempting to speculate that P19, and probably other on-switch genes expressed and secreted by neurogenic progenitors, are key triggers of microglia colonization of the brain.

While the roles of the neuro-immune crosstalk during development are receiving increasing attention, only few studies have addressed whether microglia or other macrophage types can influence the abundance and differentiation of neural progenitors as well as the specification of newborn neurons or their connectivity (Cunningham et al., 2013; Hattori and Miyata, 2018; Hattori et al., 2020; Rosin et al., 2021; Squarzoni et al., 2014). However, studies on a direct role of microglia in regulating neuronal migration are lacking. In this context, it is interesting to observe that the same on-switch gene Cxcl12 that was found to increase the density of microglia during development (Arnò et al., 2014) was also shown, in independent studies, to influence tangential migration of interneurons (Arnò et al., 2014; Borrell and Marín, 2006; Li et al., 2008). While a causal relationship linking microglia accumulation and neuronal migration by Cxcl12 is lacking, here we found that an activation and accumulation of microglia by P19 was necessary to impair neuronal migration. In turn, our study highlights a novel aspect of the neuro-immune crosstalk that is relevant in the context of neuronal migration.

On a final note, it is worth reminding that our study additionally showed that P19 can localize to the nucleus giving it potential to act beyond a secreted molecule. Assessing the functional implication of additional potential function(s) of P19 should be addressed in future studies.

## MATERIALS AND METHODS

### Bioinformatic analyses

The primary amino acid sequence of mouse P19 (UniProt ID: Q8K2W9, 541 residues) was used to infer catalytic domains by InterPro (https://www.ebi.ac.uk/interpro/) and HHPred (https://toolkit.tuebingen.mpg.de/tools/hhpred). Also, NLS Mapper (http://nls-mapper.iab.keio.ac.jp/) and NLStradamus (http://www.moseslab.csb.utoronto.ca/NLStradamus/) were used to predict nuclear localizing signals. For reference, peptide sequences with a score of 1 and 2 are predicted to localize only to the cytoplasm; scores 3-5 indicate localization in both the cytoplasm and the nucleus; scores 6 and 7 indicate partial nuclear localization; and scores 8-10 predict a mainly nuclear distribution. Additionally, TOPCONS (https://topcons.net/pred/) and Phobius (https://phobius.sbc.su.se/) were used to infer signal peptides; and PrediSi (http://www.predisi.de/index.html) and SignalP (https://services.healthtech.dtu.dk/service.php?SignalP-4.1) to infer cleavage sites in the primary amino acid sequence. Algorithms were run using default settings for eukaryotes.

### Cloning

A cDNA library was constructed using RNA extracted from embryonic mouse cortex (E14) and used as a template to clone the 4931414P19Rik gene coding sequence. To generate a P19 clone C-terminally fused with a Flag tag (P19-Flag), we employed the following primers, forward: 5’-CAACTATGTCCTTTAGTGCCA-3’; reverse: 5’-GGAAAAGGATGAATACACTCTAGACTACAAAGACGATGACGACAAGTAG -3’. This reverse primer contained the sequence of the Flag tag C-terminally fused to P19. Amplified fragments were then inserted into a pDSV-RFP backbone as previously described (Aprea et al., 2013; Artegiani et al., 2015), and validated by sequencing (Eurofins). The expression of both, P19 and RFP were under independent simian virus 40 (SV40) promoters. For the shRNAs either the ultramer 5’-GTTGACAGTGAGCGCAGGAATTATAATGCTTATCTATAGTGAAGCCACAGATGTATAGATAAGCATTATAATTCCTATGCCTACTGCCTCGGA-3’ targeting Luciferase, or the ultramer 5’-TGCTGTTGACAGTGAGCGAGGGCCAACAATGAGTTGTTAATAGTGAAGCCACAGATGTATTAACAACTCATTGTTGGCCCGTGCCTACTGCCTCGGA-3’ targeting P19 were inserted downstream the GFP sequence under the spleen-focus forming promoter (SFFV) promoter into the mir-E vector (Fellmann et al., 2013). The esiRNAs targeting either Luciferase (#RLUC) or P19 (#MU-220224-1) were purchased from Eupheria Biotech and used in combination with an empty vector expressing RFP (pDSV-RFP).

### Cell culture

HEK293cells were kept in DMEM (Thermo Fisher, cat. # 31966-047) supplemented with 10% foetal bovine serum (Gibco, cat. # 10270106) and 1% penicillin/streptomycin. Cells were used between passages 5 and 15, and regularly checked for mycoplasma contamination. The day before transfection, 4 million cells were seeded in a 10 cm petri dish. For transfection, a 3:1 mix was prepared with polyethylenimine (Sigma, cat. # 408727) and DNA, that was added to the cells. After 24 h, the transfection mix was removed and cells washed twice with PBS and replenished with 10 ml of fresh medium without serum. After additional 24 h ∼70% of the cells were transfected, and the conditioned medium and cell lysates collected. To discard detached cells and debris, the conditioned medium (10 ml) was centrifuged at 500 g for 10 min and supernatant collected (9 ml) and filtered with 0.22 µm filters at low speed (one drop per second). Filtered medium was concentrated using Amicon Ultra-15 Centrifugal Filter Unit (3 kDa cut-off; Millipore, cat. # UFC900308) at 5,000 g for 1 h. About 300 µl of concentrated conditioned medium was recovered, aliquoted and stored at –80 °C. Cells were lysed by scrapping in RIPA buffer supplemented with EDTA-free protease inhibitors (Roche, cat. # 04693159001). Lysed cells were centrifuged at maximal speed for 10 min, and supernatant collected, aliquoted and stored in the freezer until usage. See below for the use of HEK293cells on brain slices.

### Western blot

Protein lysates were denatured by heating at 70 °C for 30 min with NuPAGE LDS sample buffer (Invitrogen, cat. # NP0007) and NuPAGE reducing agent (Invitrogen, cat. # NP0004). Samples were run in 4-12% Bis-Tris protein precast gels (Invitrogen, cat. # NP0335BOX) for 1 h and 15 min at 165 V constant, using SDS running buffer MES (Invitrogen, cat. # NP0002). Proteins were transferred onto nitrocellulose membranes (0.45 µm pore size; Sigma, cat. # GE10600012) for 2.5 h at 400 mA constant, in MOPS buffer (Invitrogen, cat. # NP0001) with 15% methanol. As a reference for protein sizes, we used a pre-stained protein ladder (LI-COR, cat. # 928-60000). Immunodetections were carried out using the enhanced chemiluminescent method (Thermo Fisher, cat. # 34577).

### In utero electroporation

C57BL/6J wildtype (Janvier) mice were used, except for Supp. Fig. 1F for which the *Btg2*::GFP line was used (Haubensak 2004). The morning of the vaginal plug was defined as E0 and 13 days later (E13) mice were deeply anesthetized with isoflurane, their uterine horns exposed and approximately 2 µl of plasmid (2 µg/µl with 0.01% of Fast Green) injected into the lateral ventricle of the embryonic brains. Electrodes were placed around the embryo head, with the anode facing the injection site, and 6 pulses of 30 V for 5 ms each were delivered using an electroporator (BTX ECM830). Afterwards, the uterus was relocated within the abdominal cavity, and the muscular walls and overlying skin were sutured independently. Mice were transferred to the housing box when fully awake and sacrificed at the indicated time-points by cervical dislocation. Eventually, mice were administered with a single intraperitoneal dose of EdU (1 mg/kg) 24 h before sacrifice. All animal procedures were approved by local authorities and complied with all relevant ethical regulations (TVV 16/2018).

### Brain slice culture

Either naïve or in utero electroporated brains were dissected at E14 as described above and kept on ice-cold PBS with 0.5% penicillin/streptomycin (Gibco, cat. # 15140-122). The meninges were removed and brains immediately embedded in 4% low-melting point agarose (Carl Roth, cat. # 6351.2) and sliced using a vibratome (250 µm thick). Slices were transferred to culture inserts (Corning, cat. # 353090) pre-soaked in culture medium (Neurobasal (Thermo Fisher, cat. # 21103-049) supplemented with 10% horse serum (Thermo Fisher, cat. # 16050-130) and 1% penicillin/streptomycin. For time-lapse imaging, we labelled microglia by adding isolectin B4 conjugated to an Alexa-488 fluorophore (Thermo Fisher, cat. # I21411) to the culture medium (5 µg/ml) 2 h before imaging started. A spinning disc microscope (Andor DragonFly) was used for live-imaging at 37 °C and 5% CO_2_ with a time resolution of 15 min over a period of 12 h. For each slice, about 35 planes were acquired every 5 µm, and planes later compiled into a maximal intensity projection for analysis by ImageJ (NIH) using the Manual Tracking plugin. HEK293cells for brain slices co-culture were transfected as indicated above and two days after transfection detached and resuspended in DMEM. Collagen type I-A (Wako, cat. # 631-00651) was freshly reconstituted and mixed with the cells to a final concentration of 1.5 mg/ml. Drops of cells embedded in collagen (50 µl) were seeded on parafilm and let them solidify for 30 min at 37 °C. After solidification, the drops were placed adjacent to the pial side of naïve brain cortices, and co-cultured for 24 h. For depletion of microglia slices were incubated with liposomes (5 mg/ml) filled with PBS or clodronate (Liposoma, cat. # CP-005-005) for 48 h. At the indicated time points, the tissue was fixed with 4% PFA for 20 min, and processed for immunohistochemistry.

### Immunohistochemistry, in situ hybridization and imaging

Embryo brains were dissected and fixed overnight by immersion in 4% paraformaldehyde (PFA) and vibratome sectioning (40 µm thick) obtained. Slices were permeabilized and blocked for 1 h at room temperature in blocking buffer (PBS 0.1% Triton X-100 and 5% donkey serum) and primary antibodies (Supp. Table 2) incubated for 3 overnights at 4 °C in blocking buffer with Alexa-dye conjugated secondary antibodies (1:500, Molecular Probes) incubated for 1 overnight at 4 °C in blocking buffer. DAPI was used to counterstain nuclei and the Click-it reaction kit used to reveal EdU (Invitrogen, cat. # C10340). Immunohistochemistry was combined with in situ hybridization using the RNA-protein co-detection kit from RNAscope (cat. # 323180) following manufacturer’s instructions. P19 was detected using the Mm-4931414P19Rik-C1 probe; while the bacterial transcript dihydrodipicolinate reductase, DapB was used as negative control (PN 320871). For quantification single confocal planes were used. Tbr2 identified neuron progenitors, while apical (VZ) and basal (IZ, CP) Tbr2– cells were scored as proliferating progenitors or neurons, respectively. To reliably associate gene expression to a given cell type only discrete puncta within nuclei were scored. Images were acquired using an automated Zeiss ApoTome or confocal (LSM 780) microscope (Carl Zeiss) and maximal intensity projections quantified using ImageJ and Affinity Photo. For CD68 analysis, signal was obtained using the same settings for all the images (laser intensity, exposure time, gain, offset, etc.), and processed in ImageJ. To score CD68 mean gray intensity per microglia, the contour of Iba1+ cells were drew using the polygon selection tool, each selection was then applied to the CD68 channel and the mean gray intensity measured. Next, a threshold (40 to 255) was applied to the CD68 signal, and the binary image was used to calculate the area covered by CD68 as a percentage of the Iba1+ area.

### Statistical analyses

Quantification of cell types and morphometric analyses were performed on at least three independent biological replicates, and depicted as means ± SEM. Statistical tests were performed using Prism9 (GraphPad). Details on statistical tests and significance are indicated in the figure legends.

## Acknowledgements

We thank the facilities of the Center for Regenerative Therapies Dresden (CRTD) and Max Planck Institute of Molecular Cell Biology and Genetics (MPI-CBG) for maintenance of mouse lines, and light microscopy support. We are grateful to Wieland B Huttner for providing the *Btg2*::GFP reporter line; Mara Sannai, Francesca Bruno and Christoph Kaether for assistance with the shRNA plasmids; Michael H Sieweke for providing antibodies against CD68, CD206, and Lyve1; Simone Massalini and Melanie Jäger for excellent technical assistance, and to the members of the laboratory for critical feedback during the project.

## Author contributions

IM and FC conceived the project, analysed the data and wrote the manuscript. IM performed all experiments and quantified results.

## FIGURE LEGENDS

**Supplementary Figure 1.**
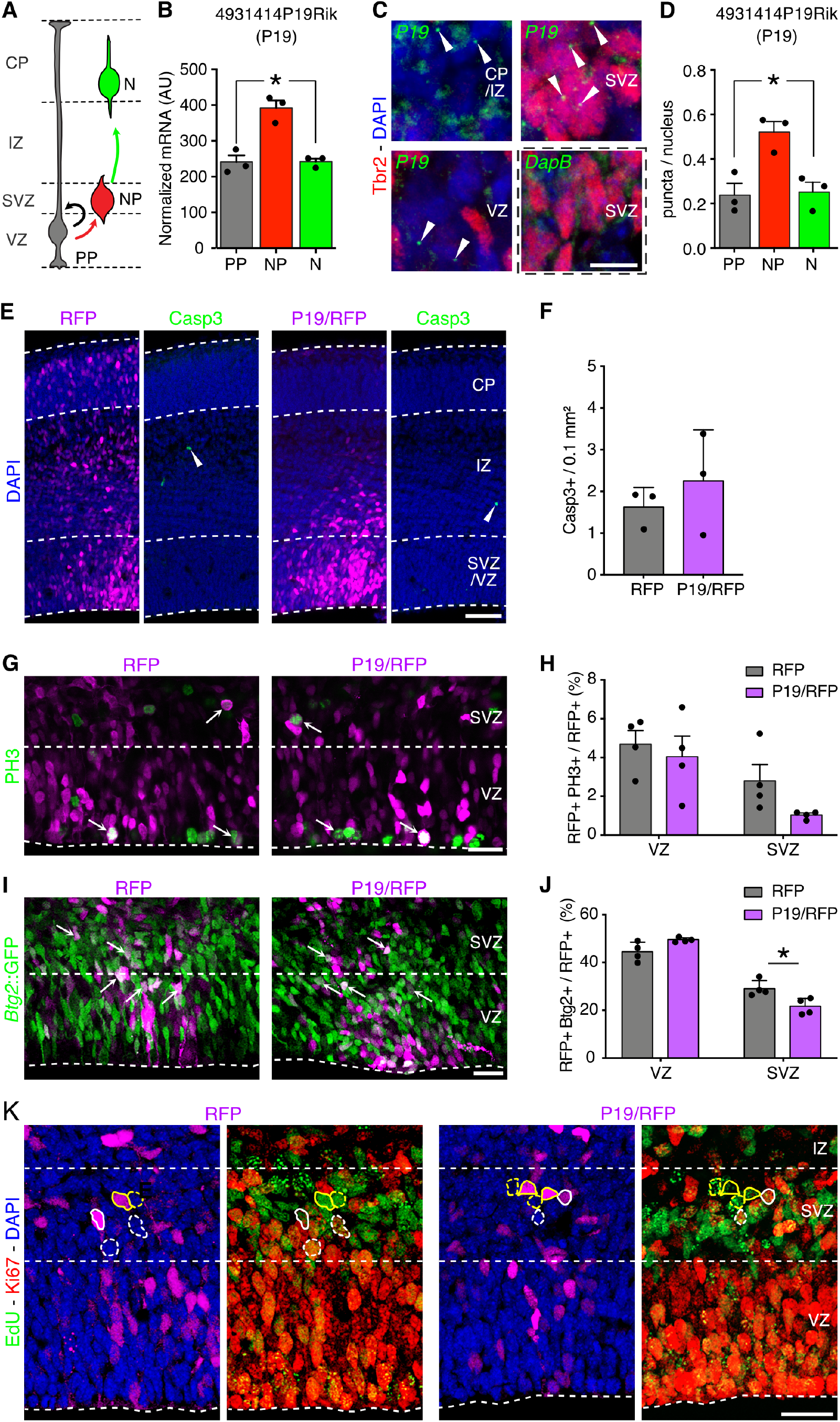
The on-switch gene P19 does not affect neural survival or mitosis. (A and B) Drawing and quantification of P19 expression levels in proliferative and neurogenic progenitors and neurons (PP, DP and N, respectively) (from Aprea et al., 2013). (C) Combined fluorescent in situ hybridization with immunolabeling in E14 coronal brain sections to detect P19 mRNA (green) and the neuron progenitor marker Tbr2 (red). Arrowheads point to discreet P19 mRNA signal. The bacterial transcript DapB was used as a negative control. (D) Quantification of P19 mRNA per cell type. (E-J) Fluorescence pictures (E, G and I) and quantifications (F, H and J) of E15 brains two days after electroporation with control RFP or P19/RFP plasmids immunolabeled with markers of apoptosis (Casp3, arrowheads), mitosis (PH3), or a reporter gene marker of neurogenic commitment (*Btg2*::GFP) as indicated. Arrows point to double positive cells. (K) Low magnification panels of those shown in Fig. 1E. Quantifications are depicted as bar graphs with individual values ± SEM. Either a Benjamini–Hochberg test (B)), one-way ANOVA and Tukey’s post-hoc test (D), a two-tailed unpaired t test (F), or a two-way ANOVA and Bonferroni’s post-hoc test (H, J) were used to assess significance (* p < 0.05). Scale bars = 10 µm (C), 50 µm (E), 25 µm (G, I, K).

**Supplementary Figure 2.**
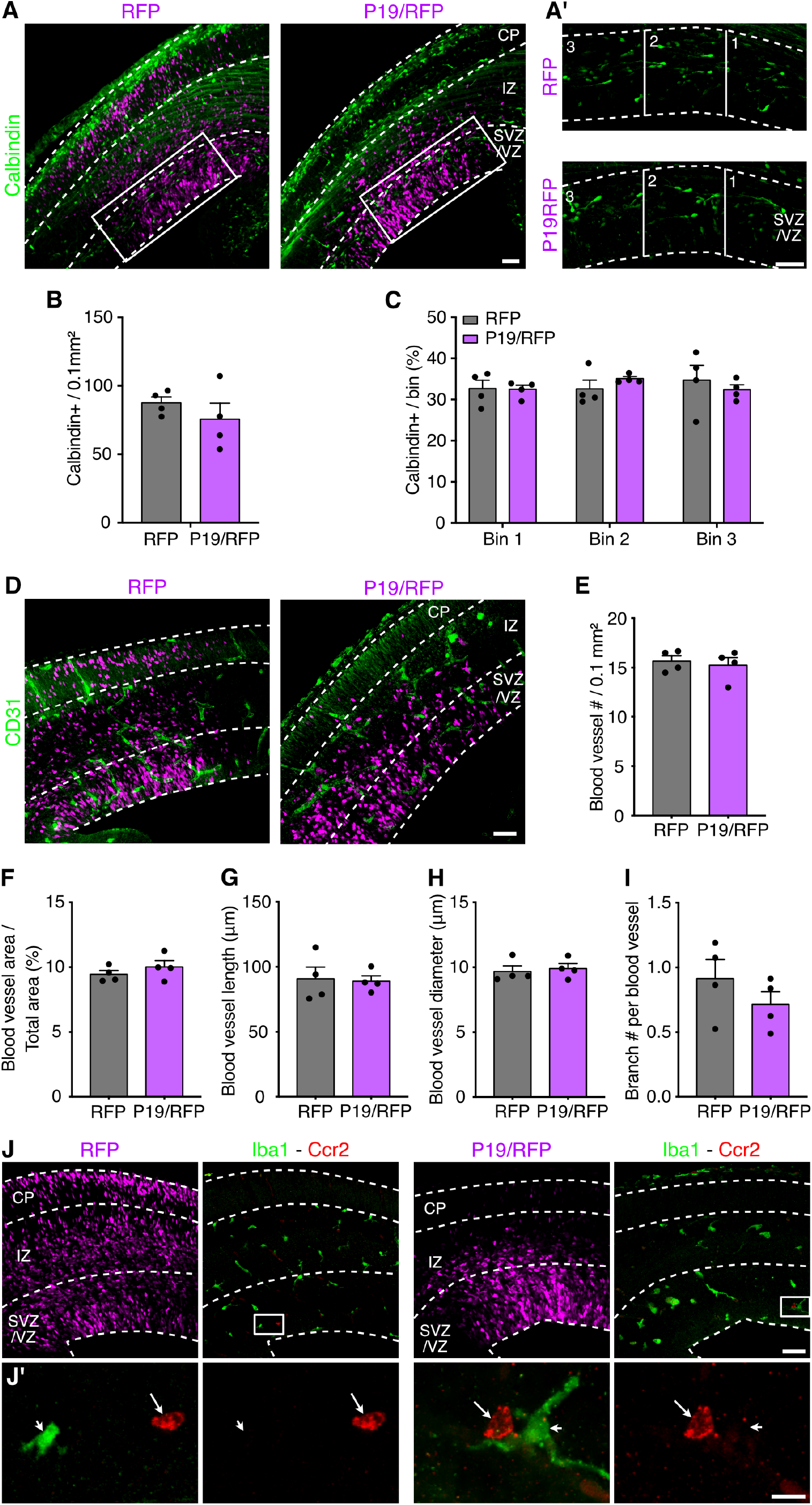
P19 overexpression does not affect interneuron migration nor angiogenesis. (A and D) Fluorescence images and corresponding quantifications (B, C, E-I) of coronal sections of E15 brains two days after electroporation with control or P19/RFP plasmids and immunolabeled with Calbindin or CD31 (as indicated) to assess interneuron migration and blood vessels architecture, respectively. Insets (A) are magnified in (A’) and continuous lines delimit bins perpendicular to the ventricular surface (A’). (J) Fluorescence images of electroporated brain sections counterstained with Iba1 (green, arrowheads) and the infiltrating macrophage marker Ccr2 (red, arrows). Quantifications are depicted as bar graphs with individual values ± SEM. Either a two-way ANOVA and Bonferroni’s post-hoc test (C), or a two-tailed unpaired t test (B, E-I) were used to assess significance. Scale bars = 50 µm (A, A’, D, J), 10 µm (J’).

**Supplementary Figure 3.**
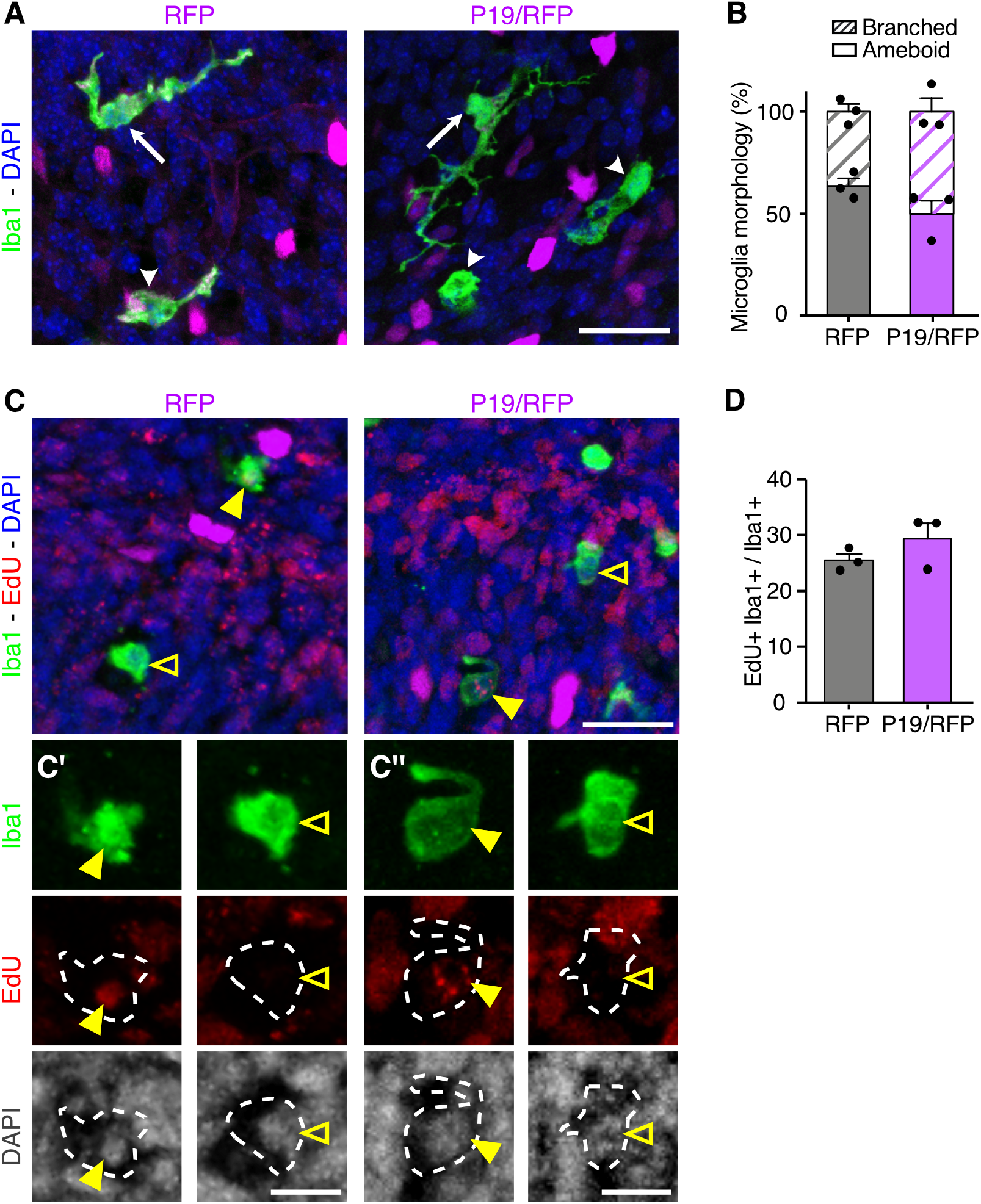
P19 overexpression does not affect microglia morphology nor proliferation. (A and C) Fluorescence images and corresponding quantifications (B, and D) of coronal sections of E15 brains two days after electroporation with control RFP or P19/RFP plasmids and immunolabeled with Iba1 and EdU (as indicated) to assess microglia morphology or proliferation, respectively. Single-channel higher magnifications are shown in (C’ and C’’). Branched or amoeboid microglia are pointed by white arrows or arrowheads, respectively. Iba1+ and EdU+ cells are pointed with a yellow arrowhead, while Iba1+ and EdU– cells are indicated with an empty arrowhead. Quantifications are depicted as bar graphs with individual values ± SEM. A two-tailed unpaired t test was used to assess significance. Scale bars = 25 µm (A, C), 10 µm (C’, C’’).

---

**Supplementary Table 1.** Predicted features of P19 amino acid sequence. Scores are given together with the maximal score or the cut-off value into parenthesis. NLS, nuclear localizing signal.

| Feature | Position | Sequence | Score | Resource |
| --- | --- | --- | --- | --- |
| NLS | 291-327 | RGTGQKNSRRKRDLVLSKLVHNVHNHITNDKRFNGS | 4.4 (max 10) | NLS Mapper |
| NLS | 369-400 | FLTKRREYRNSLNPFKGLKEKEEKKLRSRRY | 4.6 (max 10) | NLS Mapper |
| NLS | 295-302 | QKNSRRKR | > 0.3 (max 1) | NLStradamus |
| NLS | 372-402 | KRREYRNSLNPFKGLKEKEEKKLRSRRYRLF | > 0.3 (max 1) | NLStradamus |
| Signal peptide | 1-17 | MSFSATILFSPPSGSEA | 79 (max 100) | TOPCONS |
| Signal peptide | 1-17 | MSFSATILFSPPSGSEA | 0.99 (max 1) | Phobius |
| Cleavage site | 17-18 | AR | 0.62 (cut-off 0.5) | PrediSi |
| Cleavage site | 17-18 | AR | 0.52 (cut-off 0.5) | SignalP |

**Supplementary Table 2.** List of primary antibodies used in this study.

| Antibody | Host | Manufacturer | Cat. # | RRID | Dilution |
| --- | --- | --- | --- | --- | --- |
| Calbindin | rabbit | Swant | CB-38 | AB_10000340 | 1:1000 |
| Caspase3 | rabbit | BD Biosciences | 559565 | AB_397274 | 1:600 |
| Ccr2 | rabbit | Abcam | ab273050 | AB_2893307 | 1:200 |
| CD206 | rat | Biolegend | 141708 | AB_10900231 | 1:200 |
| CD31 | rabbit | Abcam | ab222783 | AB_2905525 | 1:500 |
| CD68 | mouse | Abcam | ab955 | AB_307338 | 1:500 |
| Ctip2 | rat | Abcam | ab18465 | AB_2064130 | 1:600 |
| Flag | mouse | Sigma | F1804 | AB_262044 | 1:1000 |
| GAPDH | mouse | Novus Biologicals | NB300221 | AB_10077627 | 1:1000 |
| Iba1 | rabbit | WAKO | 019-19741 | AB_839504 | 1:600 |
| Iba1 | goat | WAKO | 011-27991 | N/A | 1:400 |
| Ki67 | rabbit | Abcam | ab833 | AB_306483 | 1:500 |
| Lyve1 | rat | Invitrogen | 13-0443-82 | AB_1724157 | 1:200 |
| P2ry12 | rabbit | Abcam | ab300141 | N/A | 1:400 |
| PH3 | rat | Abcam | ab10543 | AB_2295065 | 1:600 |
| RFP | rabbit | Rockland | 600-401-379 | AB_2209751 | 1:600 |
| RFP | rat | Chromotek | 5F8 | AB_2336064 | 1:600 |
| Tbr2 | rabbit | Abcam | ab23345 | AB_778267 | 1:500 |

